# Clinical connectome fingerprints of cognitive decline

**DOI:** 10.1101/2020.10.09.332635

**Authors:** Pierpaolo Sorrentino, Rosaria Rucco, Anna Lardone, Marianna Liparoti, Emahnuel Troisi Lopez, Carlo Cavaliere, Andrea Soricelli, Viktor Jirsa, Giuseppe Sorrentino, Enrico Amico

## Abstract

Brain connectome fingerprinting is rapidly rising as a novel influential field in brain network analysis. Yet, it is still unclear whether connectivity fingerprints could be effectively used for mapping and predicting disease progression from human brain data. We hypothesize that dysregulation of brain activity in disease would reflect in worse subject identification. Hence, we propose a novel framework, *Clinical Connectome Fingerprinting*, to detect individual connectome features from clinical populations. We show that “clinical fingerprints” can map individual variations between elderly healthy subjects and patients undergoing cognitive decline in functional connectomes extracted from magnetoencephalography data. We find that identifiability is reduced in patients as compared to controls, and show that these connectivity features are predictive of the individual Mini-Mental State Examination (MMSE) score in patients. We hope that the proposed methodology can help in bridging the gap between connectivity features and biomarkers of brain dysfunction in large-scale brain networks.

## Introduction

Alzheimer’s disease (AD) is the most common form of dementia worldwide. It is well known that the pathophysiological processes start years, and possibly decades, before the clinical onset (*1*). Consequently, the identification of subjects carrying a high risk of developing the disease is necessary to study the early stage of AD pathophysiology and to adopt new and more successful therapeutic approaches. This led to the definition of the clinical construct of *mild cognitive impairment* (MCI) (*2*). According to the first conceptualization, MCI has been regarded as a clinical condition characterized by an objective memory impairment not yet encompassing the definition of dementia, but with a higher risk of developing severe cognitive decline (*2*). Currently, MCI patients are classified according to type and number of affected cognitive domains. This clinical classification is particularly relevant because each subtype is linked to a presumed etiology, in fact the amnestic subtypes (aMCI) seems to represent the prodromal form of AD (*2*).

Typically, the main symptom in aMCI is memory impairment. However, when this condition progresses toward the overt dementia phase, several cognitive functions become compromised, such as comprehension, communication, problem-solving, abstraction, imagining, planning, logic reasoning and abstract thought. To date, it has not been possible to link such functions to the malfunctioning of any specific area. This could be due to the fact that such complex abilities might not stem from a single dysfunctional area, but rather from the coordinated activity of multiple brain regions, which can be represented as a brain network or connectome (*3*).

In brain networks, nodes correspond to grey-matter regions (based on brain atlases or parcellations), while links or edges correspond to connections (either structural or functional) among them (*4*). Recent advances in functional neuroimaging have provided new tools to measure these connections, by exploiting the statistical dependencies between brain signals, giving rise to the field of functional connectivity or functional brain connectomics (*5*). Examining functional connectivity in the human brain offers unique insights on how integration and segregation of information relates to human behavior and how this organization may be altered in diseases (*6*). Indeed, considerable evidence has confirmed that anomalies in either the co-activations, the synchronization and/or the topology of the brain network are likely occurring in MCI (*7, 8*).

Despite the progress made in this direction, two main problems have arisen when using brain network models as a way to detect functional connectivity alterations in AD and/or MCI. Firstly, the clinical interpretation became more challenging, and behavioral correlates necessary to interpret the findings. Secondly, lack of replicability hindered the generalization of the results (*9*). Hence, despite considerable efforts from the community, reliably linking functional alterations to the MCI condition in a methodologically reliable and clinically valid way has been proven elusive (*10*). However, recent work on fMRI and EEG data showed that the individual connectivity does allow reliable single-subject identification in the healthy (*11*–*13*), given that good enough test-to-test reliability is provided (*14*). Nevertheless, the relationship between reliability (“connectivity fingerprinting”) and validity (associations with disease-related biomarkers) still lacks definitive answers. In other words, how does connectivity fingerprinting relate to alterations in the diseased connectomes?

Here, we introduce a methodology to test for the reliability/validity relations in clinical populations, that we named *Clinical Connectome Fingerprinting (CCF)*. The key distinction between CCF and “standard” connectome fingerprinting is that, in the CCF framework, we compare the similarity of the connectomes across test-retest sessions in patients against controls, obtaining individual similarity scores for each patient. We use these similarity scores as biomarkers for the prediction of clinical scores associated with the disease at hand. This idea is based on the consideration that the individual similarity scores obtained from CCF might provide a summary of large-scale dysregulation taking place in diseased brains. Starting from this assumption, we further hypothesised that individual alterations in the connectivity profiles, as summarized by CCF, might be associated with clinical outcomes of widespread cognitive decline, such as Mini-Mental State Examination (MMSE).

We applied the CCF technique to source-reconstructed magnetoencephalographic (MEG) data in aMCI subjects and matched healthy subjects (HS). We started with comparing the identification performance of a variety of connectivity/synchronization metrics that are commonly used to derive functional connectomes from MEG data. We selected the best performing one for further analysis, namely the phase linearity measurement (PLM) (*15*). We observed a consistent drop in connectome fingerprinting when transitioning from healthy to aMCI.Then, in order to test reliability/validity relationships in our dataset, we used intra-class correlation coefficient (ICC) to rank the edges according to their reliability in connectome fingerprinting. That is, we identified the edges that are more stable across test-retest sessions in controls, since the fingerprinting is mostly reliant on such edges. This analysis was carried out for different frequency bands separately, likely pinpointing specific circuitry (*8*). We conjectured that the same links responsible for the lack of identifiability in the MCI cohort would also be implicated in clinically observable alterations. Hence, we show that the most reliable links in the healthy (and whose reliability drops in the MCI population, as said) are indeed the ones predicting individual global cognitive impairment in patients, as measured by the Mini Mental State Examination (MMSE) scores.

## Results

We tested the Clinical Connectome Fingerprinting (CCF, Fig. 1) framework on a resting-state MEG dataset acquired from an elderly cohort of 69 subjects, 34 healthy controls and 35 affected by amnestic Mild Cognitive Impairment. The test-retest sessions were acquired during the same day, with a ∼1-minute break from each other (see Methods for details). From the initial population of 69 subjects we excluded those who: *1)* were affected by noise or *2)* did not have two test-retest sessions. This left us with 30 subjects per group, a total of 60 subjects.

**Fig. 1.**
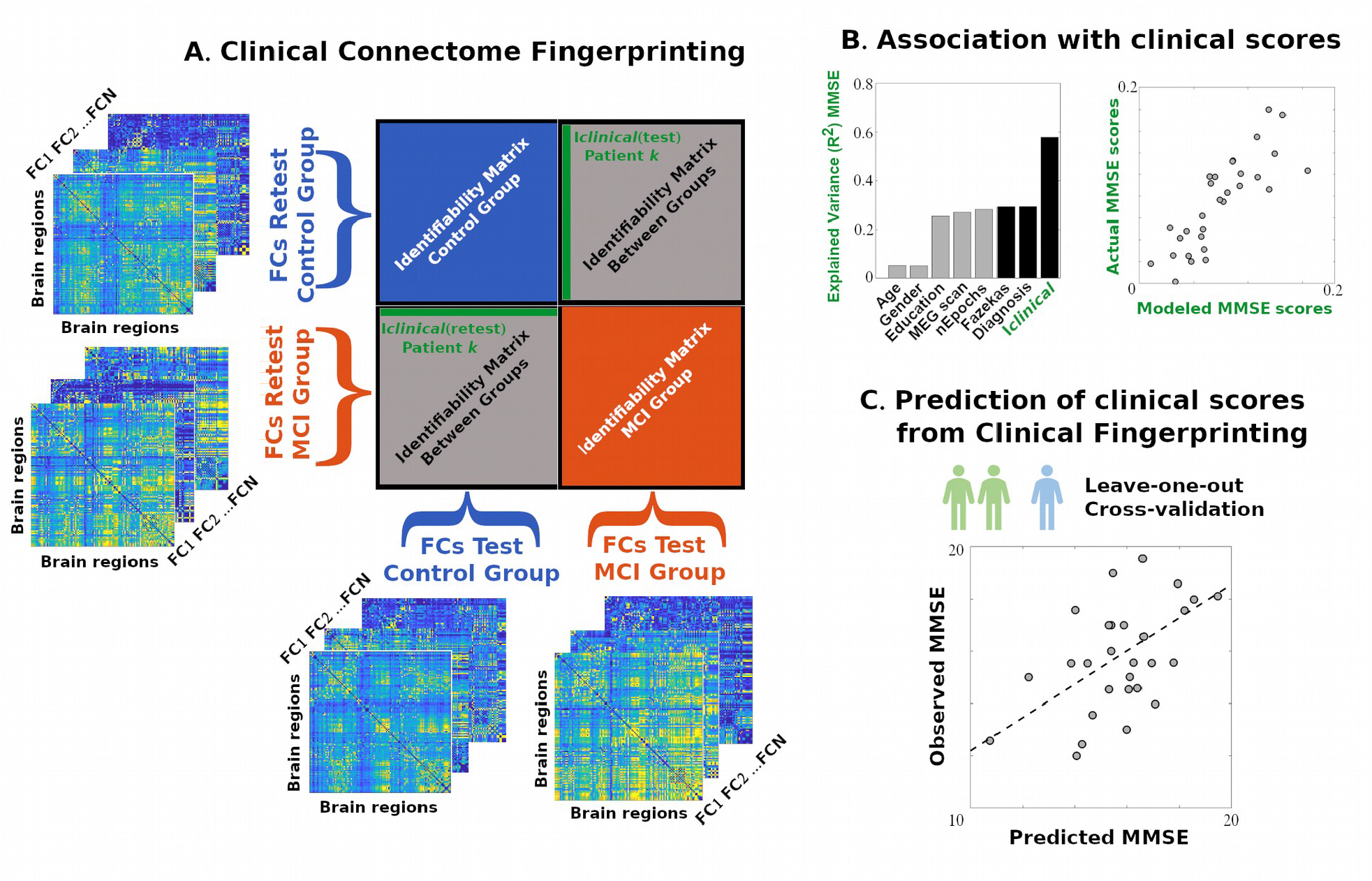
Clinical connectome fingerprinting scheme. **A)** The Identifiability (Identification) matrix (*11*) is computed for each group, using the test-retest individual connectomes; in case of two or more groups (see also (*16*)), the resulting block matrix is composed of “standard” identification matrices (red and blue blocks), plus the off-block elements which encode the individual similarity between subjects from different groups and sessions (gray blocks). Starting from this new concept one can define the “*Clinical Identifiability”* or I*clinical* for a patient *k* as the average similarity of the individual connectome of a patient with respect to the healthy control population (green row and column). Note that I *clinical* can be computed either from full individual connectomes, but also from specific targeted subnetworks (or submatrices) of interest (e.g. connectivity within visual area, etc.). **B)** One can then evaluate the association of Clinical Identifiability scores extracted from the patients’ individual connectomes with clinical scores of interest for the specific disease, using for instance a multi-linear model that accounts for several nuisance variables and predictors. **C)** Finally, the prediction and generalization power of the model can be tested by checking the performance in a leave-one out cross validation fashion.

Clinical connectome fingerprinting builds upon recent work on maximization of connectivity fingerprints in human functional connectomes (FCs) in health (*11*) and disease (*16*). The first step of CCF is to construct the “Identifiability” or “Identification” matrix (*11*), see also Fig. 1A and Methods), for the “combined” clinical and healthy population. In this case, the Identifiability matrix becomes a block matrix, where the number of blocks equals the number of groups (i.e. two in the case of this work, Fig. 1A and Methods). On one hand, each block represents identification within a specific clinical group (blue and red blocks in Fig. 1A). On the other hand, the between blocks (groups) elements (i.e. in the case of this paper, the two gray blocks in Fig. 1A) encode the similarity (or distance) between connectomes of subjects belonging to different groups (i.e., I*clinical*, see Methods for details), for both the test and retest session. In a nutshell, for each patient, I*clinical* provides the (average) score of how similar her/his connectome is with respect to the control subjects in the population, as well as across test-retest sessions (Fig. 1A). The major hypothesis behind this work is that the I *clinical* scores can be representative of the connectome degeneration associated with the disease, and therefore associated with the behavioral/clinical scores at hand (Fig. 1B, 1C).

In order to test for that, the first step was to select the best metric for fingerprinting the MEG functional connectomes. We therefore evaluated the fingerprinting capacity of six popular network metrics for MEG connectomics. Three of these were amplitude-based (Amplitude based correlation (AEC, (*17*)); AEC corrected for spatial leakage (AECc, (*18*)); Pearson correlation), and three were phase-based measurements (Phase Lag Index (PLI, (*19*)); weighted PLI (wPLI, (*20*)); Phase Linearity Measurement (PLM, (*15*))). In this regard, differential Identifiability (I*diff*, (*11*), see also Methods) provides a good score to test the robustness and reliability of each connectivity measurement across sessions.

Figure 2 shows the results of the fingerprinting test (Fig. 2): PLM seems to be the most reliable connectivity measurement among those, across all frequency bands (Fig. 2B). Interestingly, there is also a consistent drop in I*diff* scores when comparing the MCI group with the HS (Fig. 2A). We therefore selected PLM as the most robust method for the connectivity fingerprinting on this MEG dataset.

**Fig. 2.**
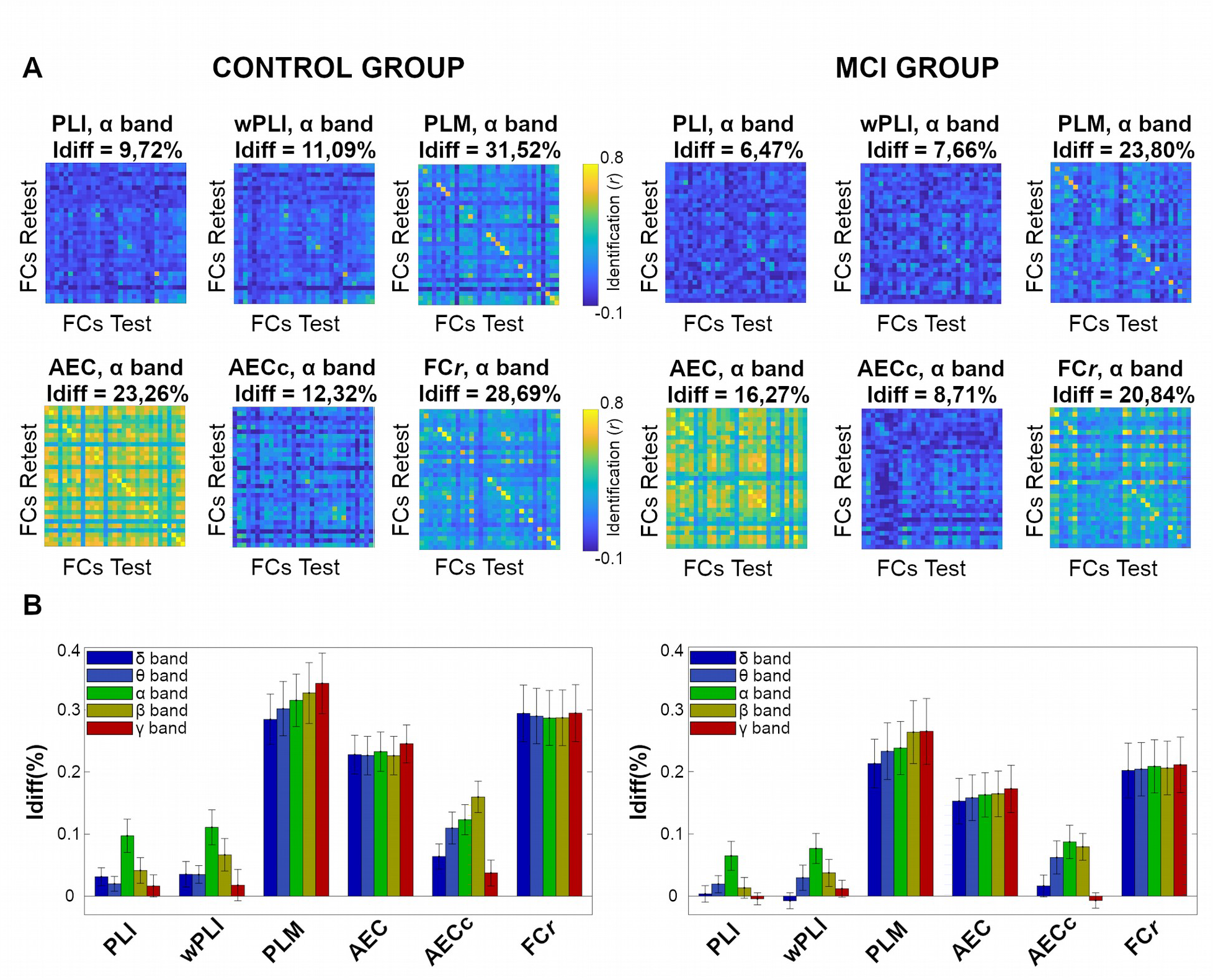
Data-driven selection of the most reliable connectivity metric for clinical connectome fingerprinting. **A)** Identifiability matrices for the HS and MCI group, for each of the six connectivity metrics tested: Phase Lag Index (PLI), weighted PLI (wPLI), Phase Linearity Measurement (PLM), Amplitude Envelope Correlation (AEC), AEC corrected for spatial leakage (AECc), Pearson’s correlation (FC*r*). Here only the alpha band is shown (the other bands are reported in Fig. S1). The differential identification score (*11*) is used to select the best metric for clinical connectome fingerprinting in this MEG dataset. **B)** The I*diff* scores across bands and connectivity metrics are summarized for the two groups; note how PLM outperforms all the other methods in all the frequency bands evaluated. We hence selected PLM connectomes for the fingerprinting analyses that follow.

We then explored the local specificity of MEG fingerprinting in PLM-based individual connectomes, by using intraclass correlation (ICC) on the functional connectome edges (see (*11*) or Methods for details), across frequency bands. Note that, in order to ease the visualization of the results, hereafter we will only show results from the three frequency bands that are most interesting to MCI, namely theta, alpha, and beta. The results for the other two (delta and gamma) are reported in SI.

The analysis on the spatial specificity of the connectome fingerprinting is depicted in Fig. 3 for the three frequencies of interest. For the control group, PLM connectivity shows the main peaks of ICC in frontoparietal and occipital regions in all bands. Somatomotor cortices show high ICC values especially in the alpha band. Note the consistent drop in ICC values when comparing the ICC edgewise patterns of the HS group to the MCI group (Fig. 3).

**Fig. 3.**
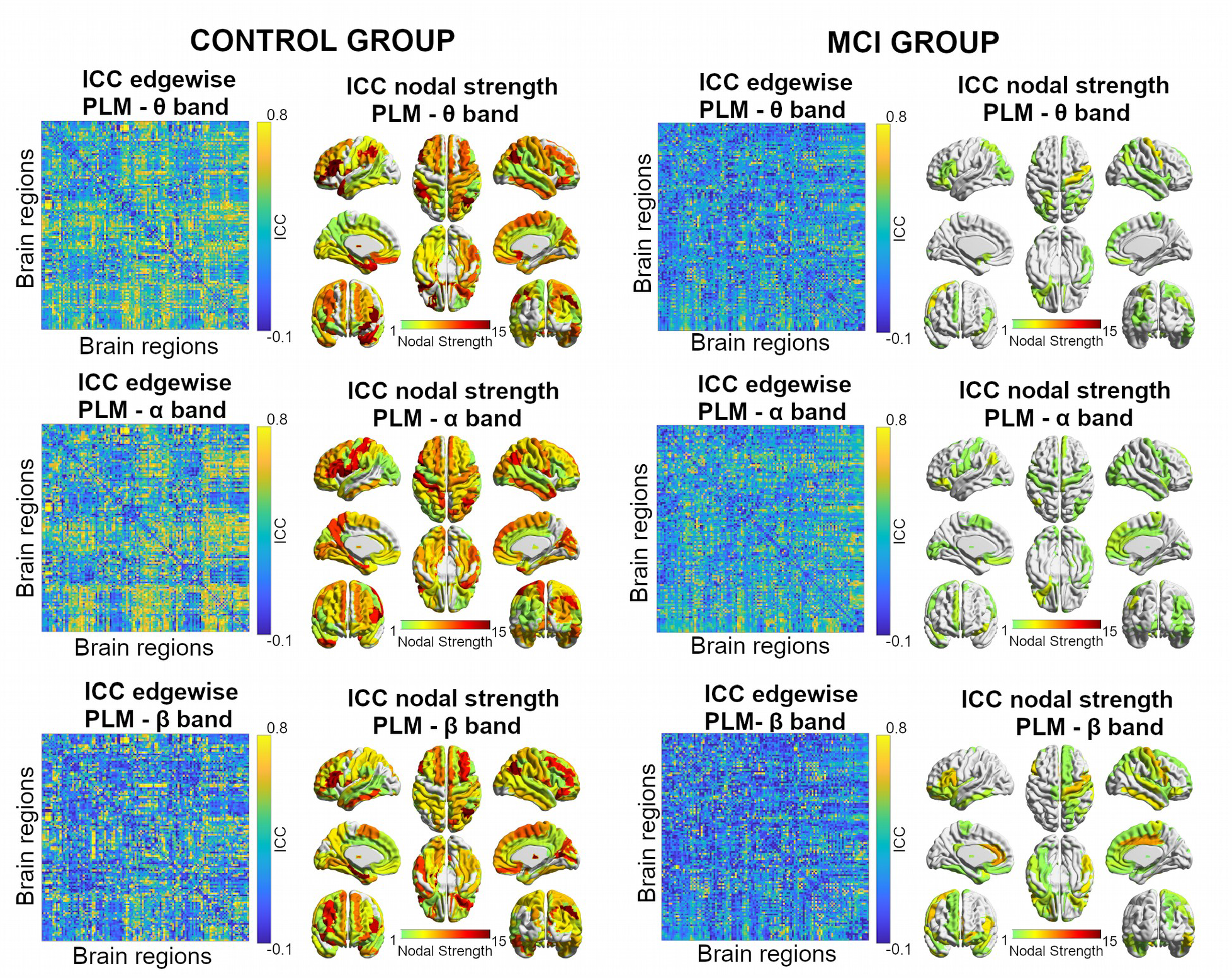
Spatial specificity of MEG connectivity fingerprints. Left: Reliability analysis of MEG connectivity fingerprints as measured via edgewise intra-class correlation (ICC), across all frequency bands (here only three are shown: theta, alpha, beta; delta and gamma are reported in Fig. S2). Right: brain renders show ICC Nodal strength of most reliable edges (greater than 75 percentile of ICC group distribution). Note the drop in the ICC distribution values when comparing the healthy control group to the MCI one.

The results reported in Fig. 3 made us speculate that a decrease in fingerprinting might be also associated with cognitive decline in the MCI population. Specifically, we sought to test the hypothesis that the individual patient’s connectome similarity/distance scores from healthy (i.e. I*clinical*, see Methods), particularly when restricted between subsets of highly reliable edges, could be used as biomarkers of cognitive decline.

We therefore tested the clinical connectome fingerprinting framework on the individual PLM matrices computed for the two groups, across all frequency bands. Briefly, we tried to predict MMSE scores from the I*clinical* similarity scores obtained from comparing connectivity subsets of most reliable edges (in an iteratively increasing fashion, from 50 to the entire functional connectomes, adding 50 edges at each iteration, analogously to (*11*)) between each MCI patient and the HS population. These I*clinical* scores were added into an additive multi-linear model to account for the possible confounds and nuisances in the dataset (Fig. 4B, see Methods for details). To test for the generalization of the prediction, similarly to Connectome Predictive Modeling (*21*), leave-one out cross validation was performed at each iteration, and the prediction score (Spearman’s ρ between predicted vs. observed MMSE, Fig. 4) was tested against the prediction score derived from two different null models: one obtained by randomly permuting the edge subset at each iteration 1000 times; the other by randomly permuting the MMSE scores 1000 times, also called permutation testing in the machine learning community (see also Methods for details). The 95% upper limit of the confidence interval obtained from permuting the MMSE labels is represented by the dashed black lines in Fig. 4A.

**Fig. 4.**
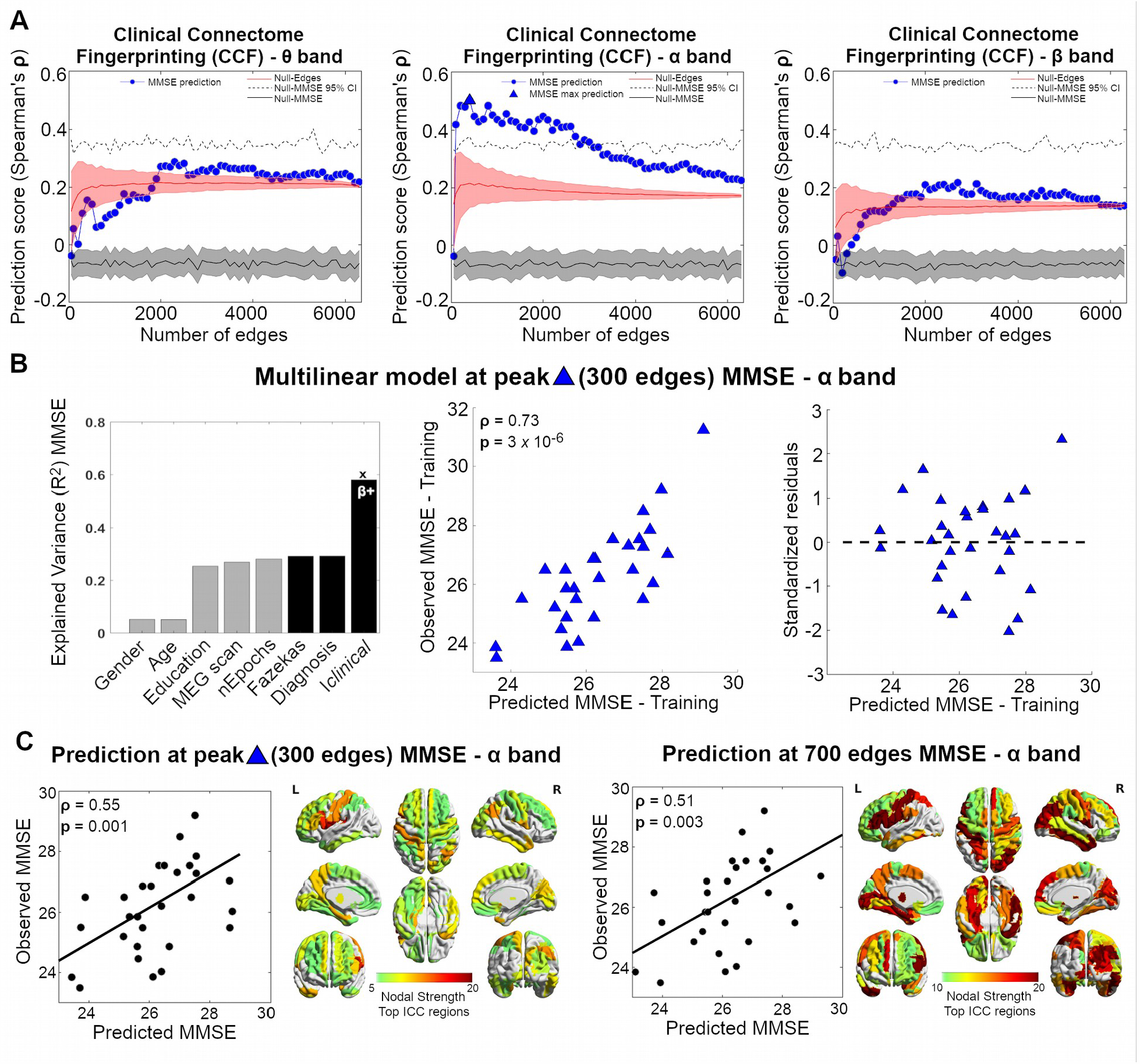
Clinical Connectome Fingerprinting for Mini Mental State Examination (MMSE) prediction. **A)** Feature selection based on ICC. At each frequency band, subset of edges are added iteratively (from 50 to whole-brain, in step of 50) based on their ICC values, from most to least reliable (x-axis), and prediction performance (Leave-one out cross validation, see Methods) of the multi-Linear model based on Clinical Identifiability (I*clinical*) is evaluated (y-axis), and compared against two null models: one (*Null-Edges*, red line), obtained by randomly choosing the subset edges 1000 times at each step (shaded red line denotes its standard deviation); the second (*Null-MMSE*, black line), obtained by randomly permuting the MMSE scores 1000 times at each step (shaded gray line indicates standard error; dashed black line denotes 95% confidence interval for *Null-MMSE*). **B)** Multi-linear model at peak. The performance of the model training set is shown for the peak prediction (300 edges, alpha band). **Left:** The additive linear model consists of five nuisance variables (Gender, Age, Education, Meg scan, number of Epochs), and three predictors (Fazekas index, Diagnosis, I*clinical*, see also Methods). Significant predictors are indicated by the **x** (p<0.05, Bonferroni corrected across frequency bands); β+ indicates that the beta coefficients for I*clinical* are positive (i.e. the higher I*clinical*, the higher the correspondent MMSE score). **Center:** Scatter plot of the Observed MMSE scores versus the MMSE scores predicted by the multi-linear regression model. **Right:** Scatter plot of the standardized residuals versus the predicted MMSE scores for the multi-linear model. Note how the residuals are symmetrically distributed, tending to cluster around 0, and within 2.5 standard deviations of zero. **C)** Nodal degree of most predictive brain regions. Figure shows prediction scatter plot for the Leave-one out test set, at peak (300 edges, alpha band) and at another local maximum (700 edges, alpha bands). The correspondent brain renders represent the nodal degree associated with the selected edge mask at 300 and 700 edges, respectively.

We found that the I*clinical*-based linear model significantly predicts the MMSE in the alpha band, with a peak in prediction when using the top 300 most reliable edges (Fig. 4A). I*clinical* scores in the training set are significantly associated with the MMSE scores (p=0.0005,R^2^*≃*0.6, Fig. 4B), with positive beta coefficients. That is, the higher I*clinical* score of the MCI patient (i.e. the more similar to the HS cohort her/his selected subnetwork), the higher her/his MMSE score. Interestingly, despite the use of a simple linear model, the LOOCV results show good generalization and prediction capacity of MMSE from connectome features, both at peak (300 edges, Spearman’s ρ=0.55, p<0.05 Bonferroni corrected across bands), or when including more edges in the selected subnetwork (e.g. 700 edges, ρ=0.51, p<0.05 Bonferroni corrected, Fig. 4C). Notably, the brain regions involved in the maximal prediction spread over the entire brain network: from frontolateral cortices, to occipital, to even cerebellar connections (Fig. 4C). Notably, prediction at alpha is significantly different from the null model based on edge permutation, as well as from the null model obtained by shuffling the MMSE scores, since the maximal prediction found is well above the confidence interval (despite the relatively small sample size, Fig. 4A).

## Discussion

In the current manuscript, we aimed to test the hypothesis that the regulation of the pattern of large-scale brain interactions is weakened in amnestic mild cognitive impairment (aMCI). Hence, we reasoned that, if the features of the functional connectome are less efficiently regulated in MCI, then the connectomes might be less easily recognizable or “identifiable”. We therefore defined a novel framework, namely *Clinical Connectome Fingerprinting* (Fig. 1), to extract individual features from diseased connectomes (or relevant subnetworks), and use them as biomarkers for prediction of cognitive decline in a MEG dataset of an aMCI population.

Firstly, we compared the identification performance of a number of commonly used MEG connectivity metrics in both the healthy and aMCI cohort. Specifically, the metrics chosen were the phase lag index (PLI) (*19*), the weighted phase lag index (wPLI), the amplitude envelope correlation (AEC) (*17*), the orthogonalized amplitude envelope correlation (AECc) (*18*), the Pearson correlation directly computed on the time series (FC*r*), and the phase linearity measurement (PLM) (*15*). Of these metrics, the FC*r*, AEC and AECc are amplitude-based, while the PLI, wPLI and PLM are phase-based. Furthermore, FCr and AEC do not correct for volume conduction, while AECc, PLI, wPLI and PLM do (although AECc uses a different approach to do so – i.e. orthogonalization (*18*)). We used source-reconstructed, resting-state MEG signals, and compared the fingerprinting capacity of the aforementioned metrics (Fig. 2). The PLM performs significantly better than the other metrics (Fig. 2). As previously known, amplitude-based metrics tend to outperform phase-metrics in terms of noise-resiliency (*22*), and metrics that do not correct for volume conduction outperform those who do in terms of identifiability (*23*), perhaps because they include information that is subject-specific though unrelated to genuine brain activity. However, PLM seems to be an exception to these trends, being a purely phase-based metric that corrects for volume conduction. One could speculate that the good resiliency of the PLM against noise (*15*) allows to extract phase information which is useful to identify subject-specific features. These features might be more related to genuine neural activity and less influenced by the geometry of the head (*22*). Importantly, provided that the identification is based on the phase of the MEG signals, this works provides complimentary information as compared to the broader fingerprint literature based on fMRI.

Once we spotted the best connectivity metric, inspired by recent work on connectome fingerprinting (*11, 16*), we tested if subject identification would be harder to perform on the MCI group as compared to controls. As shown (Fig. 3, Fig. S1), the identifiability of the patients drops drastically as compared to controls. Previous evidence showed that the large-scale activity in the healthy brain is fine-tuned to achieve both efficient communication and functional reconfiguration, which underpins complex, adaptable behavioural responses (*24, 25*). Therefore, this finding might be framed within the dysregulation of large-scale activity due to the pathological processes. Less regulated activity might imply less stable or reliable activity, which might induce lower similarity between test-retest connectomes of MCI subjects. In turn, this might imply the reduced edgewise identifiability that we observed (Fig. 3, Fig. S1).

The ICC nodal strength complements the picture of MCI altered connectivity in terms of regional activity, since not all brain regions contribute equally to subject identification (Fig. 3, Fig. S2). Interestingly, the pattern of regions that contribute the most to identifiability varies according to the frequency band. However, some constant trends appear within fronto-lateral, occipital and anterior-parietal regions (Fig. 3). It is noteworthy that some of these regions overlap consistently with previous findings in fMRI fingerprinting (*11*). However, some other areas, such as associative visual cortices (Fig. 3), which exhibit high identification power in this dataset, are not so important for fMRI fingerprinting (*12*). This led us to speculate that connectivity fingerprinting might also depend on the modality used to measure it (e.g. MEG as opposed to fMRI), which might also reflect the specific time scale of neuronal interactions/synchronies. Future studies should deepen the investigation on the relationships between fingerprinting and neuroimaging modalities.

More importantly, the main fingerprinting regions highlighted in Fig. 3 are typically associative from the functional standpoint: that is, they are believed to integrate multiple information sources and to plan coherent, complex behavioural responses (*26, 27*). We then hypothesized that impaired regulation of the interaction of such brain regions should lead to poorer cognitive performance, as well as poorer identification. If this is the case, the harder it is to identify a subject, the worse its cognitive performance should be. It is reasonable to assume that not all edges are equally important for brain fingerprinting, and that not every edge is equally affected by the pathological processes occurring in MCI. However, there might exist an overlap between these two subsets. This was indeed our working hypothesis for clinical identification.

In MCI and Alzheimer’s disease, the Mini Mental State Examination (MMSE) is one of the most widely used bedside assessments of cognitive function among the elderly (*28*). We use clinical connectome fingerprinting to show that there is a strong correlation between the MMSE score and clinical identification in the alpha band (Fig. 4A), even when taking into account confounders such as age, education, time of the acquisition, length of the scan, and subject-specific vascular burden (Fig. 4B). Furthermore, the linear model based on clinical fingerprinting scores significantly predicts the MMSE scores on leave-one out subjects (Fig. 4A, Fig. 4C). Notably, the optimal prediction is achieved when considering the first 300 most reliable edges (shown in Fig. 4). As one can observe, after including all the other covariates, the R^2^ of the model drastically increases when Clinical identifiability is taken into account (Fig. 4B). This shows that Clinical Fingerprinting might capture some processes related to cognitive performance (*14*) in MCI. Moreover, our results on Clinical Fingerprinting are obtained from a phase-based connectivity metric (PLM), which might represent specific mechanisms of communication, i.e. phase synchronization (*29*). Furthermore, the alpha band, where the best prediction occurs (Fig. 4, Fig. S3), had been previously shown to be altered in MCI (*8, 10*).

The result that the most reliable edges (high ICC (*14*)) are also predictive of an MCI-related cognitive outcome (MMSE, Fig. 4) is an interesting one. Keep in mind that, with our data-driven methodology purely based on ICC, one cannot have control over the kind of features that are being selected for the subsequent prediction of clinical outcomes (*30*). In fact, one can see that adding a few edges – despite them being the most reliable ones, as edges are being added sequentially according to their ICC – does not guarantee the best prediction. Notably however, once a sufficient number of edges has been added, one reaches the best prediction (Fig. 4A). Presumably, this prediction includes a set of edges that underpin the cognitive decline tested with MMSE. Similarly, adding further edges does not improve the prediction further, but rather makes it slowly decline (Fig. 4A). Hence, adding further (less identifiable) edges means to be adding irrelevant information for the prediction of the behavioral outcome under study. Again, our results in this MCI cohort show that the most reliable edges are also the most predictive ones, clinically. If these edges were not specifically related to the cognitive output, then a randomly selected subset of edges should perform similarly in terms of predictive power on the MMSE score. This does not seem to be the case, as the null model results show (Fig. 4A, red shaded line). In fact, when considering random edge selections, the quality of the prediction drops drastically, showing that the ICC selects those edges that are informative with respect to cognitive performance, as tested through MMSE (Fig. 4A, red shaded line).

The findings of this study make it essential to lay out several methodological considerations. The first one relates to the reliability/validity “dichotomy” (*14*). Here we show that edges that are most reliable possess a strong clinical validity for cognitive impairment prediction (Fig. 4). However, the reader should keep in mind that robust edges in MEG data can be associated with several factors, not all of them necessarily neuronal-related: motion artifacts, gray matter atrophy, individual source reconstruction parameters, epochs length, and so on. Despite our efforts in controlling for all these (as detailed in the Methods and shown in Fig. 4B), further studies should dig into the fingerprinting “causes” and properties of MEG data. The same applies to the clinical validity part of our findings: the link found between reliability/validity might be dataset and/or disease dependent, and should be explored in different populations and clinical conditions. Also, here we use a data-driven method to select the best edge features for clinical prediction, as they turn out to be significantly better than random selected features (Fig. 4A). Nevertheless, we encourage further work to explore edge selection based on a-priori hypothesis for the disease at hand, which might outperform the proposed data-driven feature selection for clinical connectome fingerprinting. Future studies should also explore the effect of denoising techniques to maximize fingerprints from brain data onto the I*clinical* scores (*11*).

Another important caveat of this study is that, in MEG source-reconstructed data, the signal-to-noise ratio is heavily dependent on the depth distance between the source and the sensor, and hence is not homogeneous for all the sources. However, recent evidence showed that signals reconstructed from the basal ganglia contain reliable information about brain activity (*31, 32*) as well as those from the cerebellum (*33*). On the one hand, MEG signals derived from the cerebellum are 30-to-60 % weaker as compared to the cortical surface (*34*). On the other hand, the cerebellum is an important structure in both motor and cognitive processes, and hence excluding it all along is likely discarding some information (*35*). In this work we tried to account for this by excluding entirely within-cerebellar links from ICC edge selection, while keeping edges that are incident but not contained within the cerebellum (see Methods for details). However, we observed that our predictions are maintained when including within-cerebellar links, while they drastically drop when excluding the cerebellum altogether from the individual connectomes (MMSE maximal prediction ρ=0.3, Fig. S4). One should keep in mind that our predictions involve a clinical outcome. The fact that cortical-cerebellar links are needed to improve MMSE prediction, after controlling for all nuisance factors and after benchmarking it against an appropriate null-model for edge selection, implies that they contain relevant information associated with the individual cognitive decline that should not be discarded. We encourage further research to avoid the underestimation of cerebellar connectivity in MCI and AD, and explore further the role of these cortico-cerebellar pathways.

In conclusion, we have defined *Clinical Connectome Fingerprinting*, a novel approach to extract individual connectivity features from diseased functional connectomes. We applied this framework for clinical identification of MEG connectomes extracted from an elderly population of subjects undergoing cognitive decline (i.e., amnestic MCI). We showed that the most identifiable edges are also the most predictive of individual cognitive impairment, as measured by the Mini-Mental State Examination score. We hope that future studies will exploit further the potential of Clinical Connectome Fingerprinting as a preclinical diagnostic tool, as well as a way to empirically link, in a data-driven fashion, specific sub-networks to given cognitive functions or brain states.

## Materials and Methods

### Participants

For this study, eighty-six patients referring to the Center for Cognitive and Memory Disorders of Hermitage Capodimonte Clinic in Naples were consecutively recruited. All subjects, aged 53 to 81, were right handed (none of them had any left-handed relatives) and native Italian speakers. Thirty-four age-gender-body mass index (BMI)- and education-matched subjects among patients spouses or friends were enrolled as control group (HSs). Exclusion criteria were the presence of neurological or systemic illness that could affect the cognitive status, and contraindications to MRI or MEG recording.

Both patients and HS underwent the following screening: neurological examination, extensive neuropsychological assessment (see Table 1), MRI scan (including hippocampal volume evaluation) and MEG recording. MCI diagnosis was formulated according to the National Institute on Aging-Alzheimer Association (NIA-AA) criteria (*36*), which include: (i) cognitive concern reported by patient or informant or clinician, (ii) objective evidence of impairment in one or more cognitive domains, typically including memory, (iii) preservation of independence in functional abilities, (iv) not demented. Reduced hippocampal volume detected by structural MRI, gives our aMCI cohort an intermediate likelihood of being due to AD (*36*).

**Table 1.** Neuropsychological evaluation. MMSE: Mini Mental State Examination (*28*); FAB: Frontal Assessment Battery (*37*);; BDI: Beck Depression Inventory (*38*); MDB: Mental Deterioration Battery (*39*); FCSRT: Free and Cued Selective Reminding Test (*40*).

| Test | Explored function |
| --- | --- |
| MMSE | Global cognitive status |
| FAB | Frontal efficiency |
| BDI | Depression |
| MDB |  |
| Rey's 15 word immediate recall | Short and long-term verbal episodic memory |
| Rey's 15 word delayed recall |  |
| Word fluency | Ability to access lexical-semantic memory store |
| Phrase construction | Language |
| Raven's 47 progressive matrices | Conceptual reasoning |
| Immediate visual memory | Short-term visuo-perceptual recognition memory |
| Freehand copying of drawings | Constructive apraxia |
| Copying drawings with landmarks |  |
| FCSRT |  |
| FCSRT immediate free recall | "Hippocampal" episodic memory |
| FCSRT immediate total recall |  |
| FCSRT delayed free recall |  |
| FCSRT delayed total recall |  |
| FCSRT index of sensitivity of cueing |  |

Screened subjects with either MRI alterations (traumatic brain injury, meningioma, lacunar infarction), diagnosis of depression, dementia or non-amnestic MCI were excluded from further analysis. The subjects included in the study were 35 patients affected by aMCI (mean±SD age 71.20±6.67 years; 18 men and 17 women) compared to 34 age, educational level and gender matched healthy subjects (mean±SD age 69.88±5.56 years; 19 men and 15 women).

The study was approved by the Local Ethics Committee “Comitato Etico Campania Centro” (Prot.n.93C.E./Reg. n.14-17OSS), and all subjects gave written informed consent. All methods included in the protocol were carried out in accordance with the Declaration of Helsinki.

### Magnetic Resonance Imaging acquisition

For 24 HS and 32 MCI patients, MR images were acquired using a 3T Biograph mMR tomograph (Siemens Healthcare, Erlangen, Germany) equipped with a 12 channels head coil. The scan was performed after the MEG registration or at least 21 days before (within 1 month). The MR registration protocol was: (i) three-dimensional T1-weighted Magnetization-Prepared Rapid Acquisition Gradient-Echo sequence (MPRAGE, 240 sagittal planes, 214 × 21 mm2 Field of View, voxel size 1 × 1 × 1 mm3, TR/TE/TI 2,400/2.5/1,000 ms, flip angle 8°); (ii) Three-dimensional T2-weighted Sampling Perfection with Application optimized. Contrasts using different flip angle Evolution sequence (SPACE, 240 sagittal planes, 214 × 214 mm2 Field of View, voxel size 1 × 1 × 1 mm3, TR/TE 3,370/563); (iii) Two-dimensional T2-weighted turbo spin echo Fluid Attenuated Inversion Recovery sequence (FLAIR, 44 axial planes, 230 × 230 mm2 Field of View, voxel size 0.9 × 0.9 × 0.9 mm3, TR/TE/TI 9,000/95/25,00, flip angle 150○). The volumetric analysis was performed using the Freesurfer software (version 6.0) (*41*), specifically the normalization of the volumes was made by the estimated total intracranial volume (eTIV) while the Fazekas scale was used to evaluate the vascular burden (*42*). For the remaining participants who refuse or did not complete the MR scan we used a standard MRI model.

### MEG Acquisition and Preprocessing

The data were acquired using a MEG system equipped by 163 magnetometers SQUID (Superconducting Quantum Interference Device) (*43*). 154 of them are located to be as close as possible to the head of the subjects, the remaining, organized into three triplets, are positioned more distant from the helmet to measure the environmental noise.

MEG data were acquired during two eyes-closed resting state segments, each 3.5 minutes long, with a minute distance between them. During the acquisition, subjects were seated inside a magnetically shielded room (AtB Biomag, Ulm, Germany) in order to reduce the external noise. Using Fastrak (Polhemus^®^) we digitalized the position of four anatomical landmarks (nasion, right and left pre-auricular points and vertex of the head) and the position of four reference coils (attached to the head of the subject), in order to define the right positions of the head under the helmet. The coils were activated and the position of the head was checked before each segment of registration. During the acquisition, we recorded also the cardiac activity and the eyes movements in order to remove physiological artefacts. After an anti-aliasing filter, the data were sampled at 1024 Hz.

The MEG data were filtered in the band 0.5-48 Hz using a 4th-order Butterworth IIR band-pass filter, implemented offline using Matlab scripts within the Fieldtrip toolbox (*44*). As described previously (*45*), Principal Component Analysis was applied to reference SQUID signals to remove the environment noise. Subsequently, noisy channels and bad segments of acquisition were identified and removed through visual inspection by an experienced rater. Finally we removed physiological artifacts, such as eye blinking and heart activity, by means of Independent Component Analysis. 5 MCI patients and 4 HS were excluded due to their low-quality recordings.

### Source reconstruction

Firstly, to reconstruct time series related to the centroids of 116 regions-of-interest (ROIs), derived from the Automated Anatomical Labeling (AAL) atlas, we used Nolte’s volume conduction model and the Linearly Constrained Minimum Variance (LCMV) beamformer algorithm (for details see (*46*)), based on the native MRIs. Then, we filtered the time series in the five classical frequency bands (delta (0.5 - 4.0 Hz), theta (4.0 - 8.0 Hz), alpha (8.0 – 13.0 Hz), beta (13.0 – 30.0 Hz) and gamma (30.0 – 48.0 Hz)). Figure 5 shows the data analysis pipeline.

**Figure 5.**
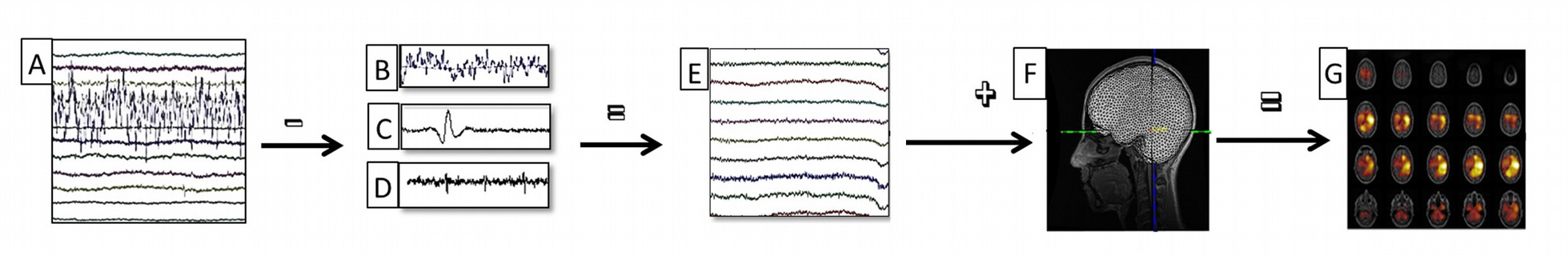
Data analysis pipeline. (A) Raw MEG signals recorded by 154 sensors (a subset displayed here). (B-C-D) Respectively noisy channel, cardiac artifact, blinking artifact, removed during preprocessing phase. (E) MEG signals after noise cleaning and artifact removal. (F) Co-registration between MEG signals and MRI. (G) Source reconstruction (beamforming).

### Functional connectivity measurements

As connectivity measurements we used three amplitude-based and three phase-based metrics. Specifically, as amplitude-based metrics we used i) the classical functional connectivity based on the Pearson’s correlation between brain signals (FCr); ii) Amplitude envelope correlation (AEC) (*17*) which computes the amplitude envelope by means of the Hilbert transform and then determines functional connectivity between brain signals through the Pearson correlation coefficient; iii) the orthogonalized Amplitude Envelope Correlations (AECc) with signal leakage correction (*18*). As phase-based metrics we considered i) the Phase Lag Index (PLI) which estimates the asymmetry of the distribution of the phase differences between the brain signals (*19*); ii) the weighted Phase Lag Index (wPLI) which weights the PLI by the magnitude of the imaginary component of the cross-spectrum (*20*); iii) the Phase Linearity Measurement (PLM) which measures the synchronization between brain regions by monitoring their phase differences in time (*15*).In conclusion, for each subject and each metric, we obtained two-test retest connectomes.

### Towards Clinical Connectome Fingerprinting

The methodology for clinical connectome fingerprinting is inspired by recent work on maximization of connectivity fingerprints in human functional connectomes in health (*11*) and disease (*16*). Briefly, it starts from defining the mathematical object known as “Identifiability” or “Identification” matrix (*11*) see also Fig. 1A).The identifiability matrix has subjects as rows and columns, and encodes the information about the self similarity (I*self*, main diagonal elements) of each subject with herself/himself, across the test/retest sessions, and the similarity of each subject with the others (or I *others*, off diagonal elements) (*11*). In this context, the similarity is defined as the Pearson’s correlation between the connectomes at hand. The difference between the average I*self* and the average I*others* (denominated *“Differential Identifiability”* or *“Differential Identification” -* I*diff* (*11*)) provides a robust score of the fingerprinting level of a specific dataset (*11*).

This framework can easily be extended in scenarios where multiple clinical groups are present ((*16*) see also Fig. 1A). In this case, the Identifiability matrix becomes a block matrix, where the number of blocks equals the number of groups (i.e. two in the case of this work, Fig. 1A). The within-group blocks (blue and red blocks in Fig. 1A) represent the Identifiability matrix *within* a specific clinical group (i.e. MCI or healthy controls). The between blocks (groups) elements (i.e. the two gray blocks in Fig. 1A) encode the similarity (or distance) *between* the test-retest connectomes of subjects belonging to different groups. In particular, the top right block contains the similarities between the connectomes from patients during the test session with the connectomes of the controls during the retest session, while the bottom-left block contains the opposite case (i.e. the similarities between the connectomes of the patients during the re-test session with the connectomes of the controls during the test session). Let *C* be the set of the healthy volunteers. Similarly, let *G* define the patient group. Also, let I be the Identifiability matrix depicted in Fig. 1A. Hence, we can define I*clinical*(test) and I *clinical* (retest), for a specific patient *k*, as:

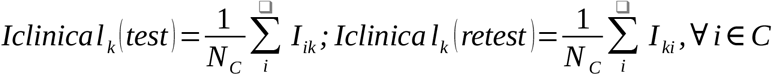

In a nutshell, *Iclinical(test), for each patient k*, represents the average similarity of the connectome of that patient in the test session with the connectomes of every control in the retest session, and I*clinical*(retest) represents the average similarity of the connectome of a patient in the retest session with the connectomes of every control in the test session. Taking advantage of the new piece of information provided by the between groups blocks, we define the “Clinical Identification” or “Clinical Identifiability” (I*clinical), for patient k*, as:

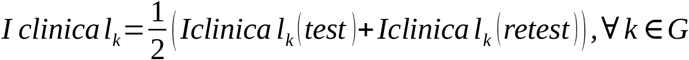

In summary, for each patient *k*, I*clinical* provides the (average) score of how similar is her/his connectome with respect to the control subjects in the population, across the test-retest sessions.

### From Clinical Connectome Fingerprinting to prediction of clinical scores

#### Edgewise Intraclass correlation

For each group (HS and MCI) we quantified the edgewise reliability of individual connectomes using intraclass correlation (ICC, (*47*)), similarly to previous work (*11*). ICC is a widely used measure in statistics, normally to assess the percent of agreement between units of different groups. It describes how strongly units in the same group resemble each other. The stronger the agreement, the higher its ICC value. We used ICC to quantify to which extent the connectivity value of each edge (functional connectivity value between two brain regions) could separate within and between subjects. In other words, the higher the ICC of an edge, the more stable that edge’s connectivity across the test-retest session and, in turn, the higher the edge’s “fingerprint” (i.e. that particular edge is relevant to identify individual connectomes). We then go on and test if the edge’s identifiability diminishes in patients, and if such reduction is predictive of individual clinical disability.

#### Multilinear model specification

To test for the hypothesis that clinical connectome identification is associated with clinical scores, we performed a multi-linear regression analysis to predict the MMSE scores based on I*clinical* and two other predictors. Specifically, a categorical variable encoding diagnosis (amnestic MCI and multi domain MCI) membership, and the Fazekas index which quantifies the amount of white matter T2 hyperintense lesions (i.e. vascular burden). Five nuisance variables were also included to account for any potential effects of age, sex, education, different day of MEG scans, and different number of epochs.

#### Edge selection and prediction of clinical scores

The Clinical Identification scores defined earlier (Fig. 1A) can be computed from full individual connectomes, but also on a subset of the individual connectomes (i.e. by computing Patient/Controls similarity only on a subset of edges). Furthermore, if the reduction of identifiability is related to the pathological processes, we expect that the individual level of fingerprinting would be predictive of the individual clinical impairment, and maximally so when based on the subset of most reliable edges. Therefore, we tested the specificity of the prediction, as well as the generalization capacity of our model, by using a leave-one out cross validation (LOOCV) approach. The approach detailed below has some similarities with the Connectome Predictive Modeling methodology (CPM, (*48*)), with two major key differences.

Firstly, we selected the connectome edges to be included in the fingerprinting based on the edgewise ICC value computed on the control group. That is, edges were ranked in descending order according to the ICC value, and only a subset was included in the fingerprint analysis (similarly to (*11*)). Note that, in order to improve the signal-to-noise ratio and avoid source reconstruction artefacts, edges entirely within the cerebellum were not included in the analysis. LOOCV was then performed iteratively by adding 50 edges at the time, starting from the most reliable edges (as measured by ICC), and ending with the least robust ones, until eventually taking into consideration the full individual connectomes. Secondly, as aforementioned, at each iteration the individual I*clinical* scores, based on the iteration-specific subset of edges, were used to predict the individual MMSE scores. As said, depending on the number of edges included, the I*clinical* represents the similarity with (or distance from) the control group relative to the specific connectome subcircuit spanned by the included edges. Finally, the prediction scores between the ML model with I*clinical* and MMSE clinical scores were evaluated for each of the five frequency bands studied.

#### Null models for prediction

In order to make sure that the edge selection based on the ICC scores is clinically meaningful, we implement two different null models. In the first one (named *Null-Edges*), we built a distribution of the prediction scores based on randomly selected edges. Hence, at each step of the LOOCV model we shuffled the ICC mask 1000 times, and recomputed the prediction scores between the Iclinical multi-linear model and MMSE. In other words, we build a “null distribution” of prediction rates, entirely based on a randomized edge selection, however starting from the empirical functional connectomes. In the second null model, we are accounting for the sample size effect when performing LOOCV. LOOCV has been shown – both theoretically and empirically – to lead to unrealistic and unstable prediction estimates in small neuroimaging datasets (*49, 50*). Permutation testing is one possible way to account for bias in prediction estimates (*49, 50*). Hence, in the second null model (named *Null-MMSE*) we performed permutation testing on the MMSE scores, by randomly shuffling them 1000 times, and by computing the corresponding confidence intervals related to the obtained null distribution. Prediction scores outside of the 95 percentile of the Null-MMSE distribution were considered as significant predictions for our model.

## Acknowledgments

EA acknowledges financial support from the SNSF Ambizione project “Fingerprinting the brain: network science to extract features of cognition, behavior and dysfunction” (grant number PZ00P2_185716).

## Data availability

The data that support the findings of this study are available on request from the corresponding authors. The data are not publicly available due to them containing information that could compromise research participant privacy/consent.

## Code availability

The code (in MATLAB) used for this analysis will be available upon acceptance on EA EPFL webpage.

## Author Contributions

PS and RR collected and acquired the dataset, processed the data and conceptualized the study; EA conceptualized the study, designed the framework and performed the connectivity analyses; all authors interpreted the results and wrote the manuscript.

