## Supplementary Information for "Clinical connectome fingerprints of cognitive decline"

### Supplementary Materials

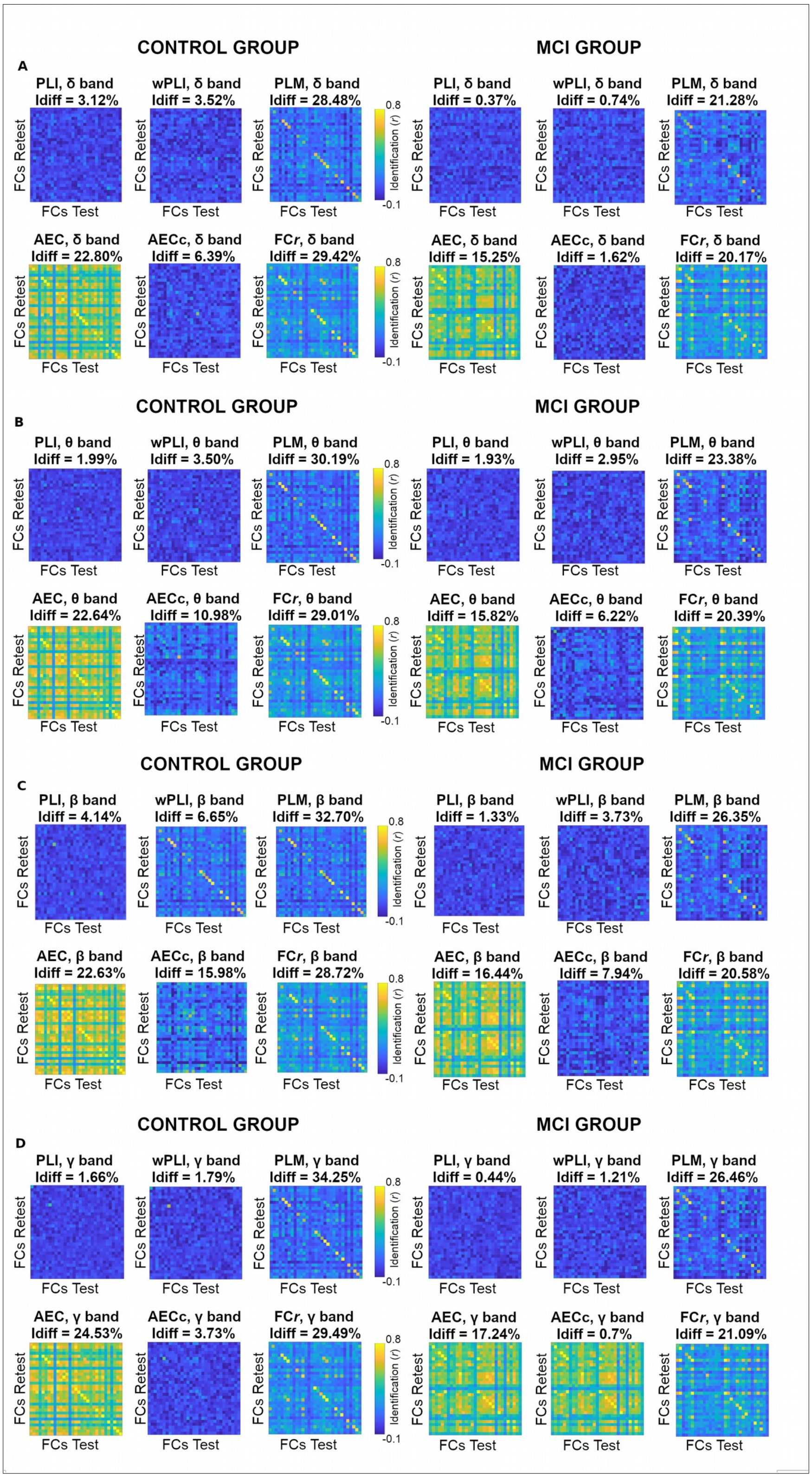

**S1 Data-driven selection of the most reliable connectivity metric for clinical connectome fingerprinting.** Identifiability matrices for the HS and MCI group, for each of the six connectivity metrics tested, for delta, theta, beta and gamma bands. The differential identification score (Idiff) is used to select the best metric for clinical connectome fingerprinting in this MEG dataset.

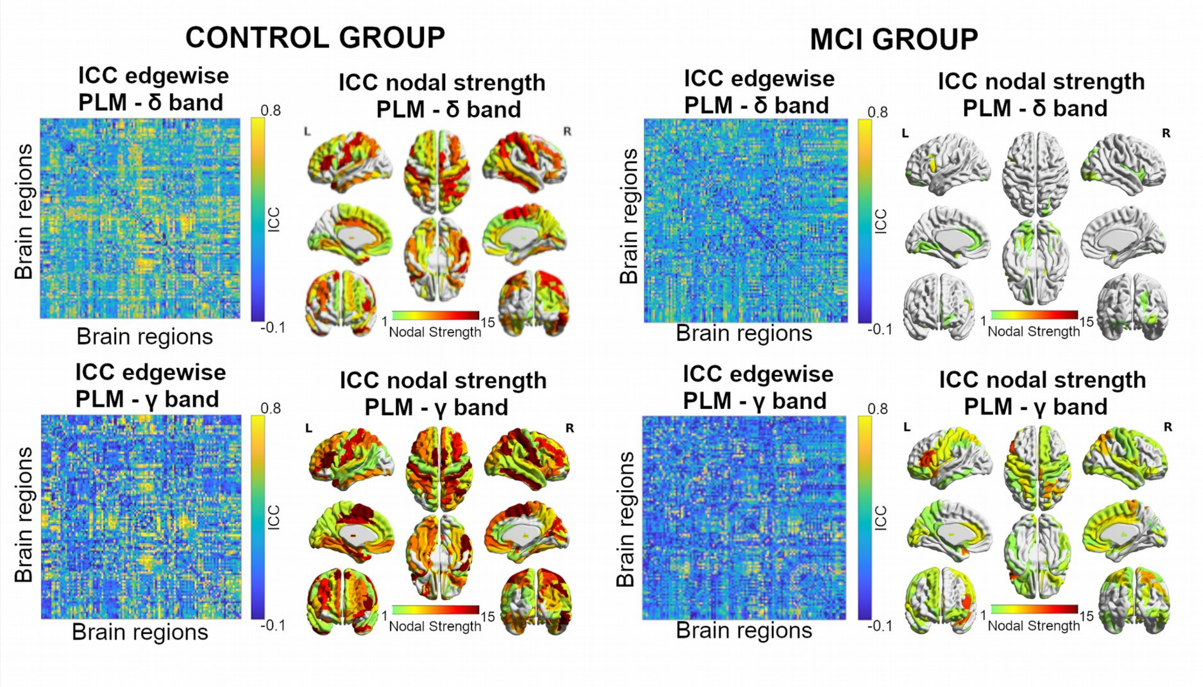


**S2 Spatial specificity of MEG connectivity fingerprints.** Left: Reliability analysis of MEG connectivity fingerprints as measured via edgewise intra-class correlation (ICC), in delta and gamma bands, for both HS and MCI groups. Right: brain renders show ICC Nodal strength of most reliable edges (greater than 75 percentile of ICC group distribution). Note the drop in the ICC distribution values when comparing the healthy subjects group to the MCI one.

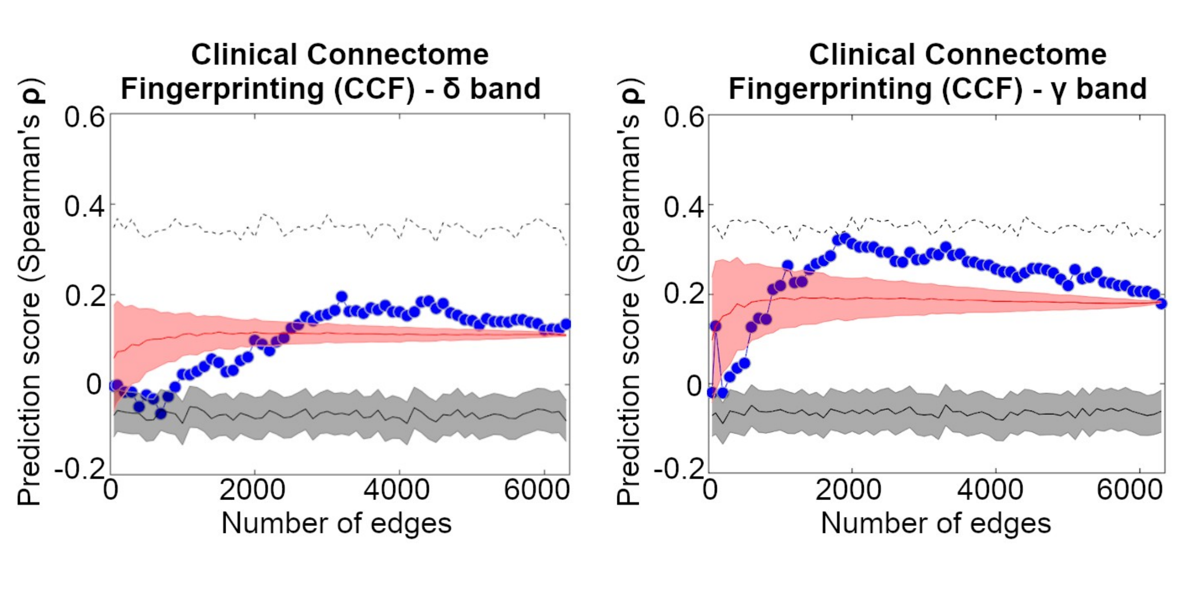


**S3 Clinical Connectome Fingerprinting for Mini Mental State Examination (MMSE) prediction.** Feature selection based on ICC, in delta and gamma bands. At each frequency band, subset of edges are added iteratively (from 50 to whole-brain, in step of 50) based on their ICC values, from most to least reliable (x-axis, see also Fig. 2), and prediction performance (Leave-one out cross validation, see Methods) of the multi-Linear model based on Clinical Identifiability (I*clinical*) is evaluated (y-axis), and compared against two null models: one (*Null-Edges,* red line), obtained by randomly choosing the subset edges 1000 times at each step (shaded red line denotes its standard deviation); the second (*Null-MMSE,* black line), obtained by randomly permuting the MMSE scores 1000 times at each step (shaded gray line indicates standard error; dashed black line denotes 95% confidence interval for *Null-MMSE*).

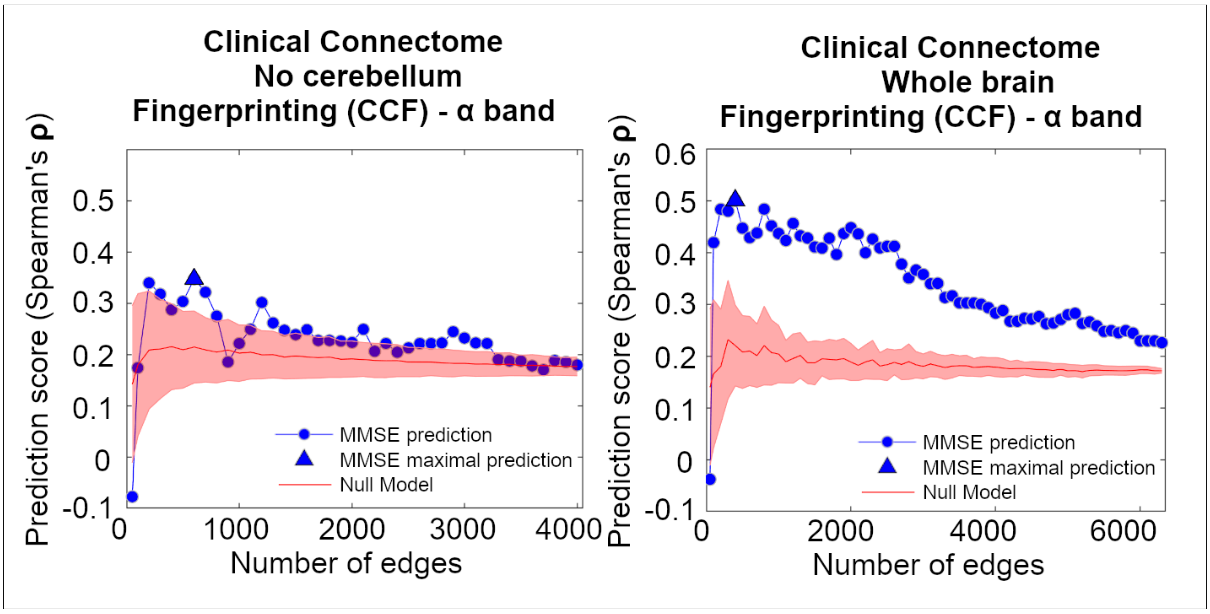


**S4 Clinical Connectome Fingerprinting for Mini Mental State Examination (MMSE) prediction.** Feature selection based on ICC, in alpha band. Right is whole brain, left is without considering cerebellum. At each frequency band, subset of edges are added iteratively (from 50 to whole-brain, in step of 50) based on their ICC values, from most to least reliable (x-axis, see also Fig. 2), and prediction performance (Leave-one out cross validation, see Methods) of the multi-Linear model based on Clinical Identifiability (I*clinical*) is evaluated (y-axis), and compared against a null model obtained by randomly choosing the subset edges 1000 times at each step (shaded red line denotes one standard deviation).
